# Investigating the active substance and mechanism of Jing-Fu-Kang granules via mass spectrometry technology and network pharmacology method

**DOI:** 10.1101/2021.08.09.455734

**Authors:** Xin Feng, Yuelin Bi, Xuhua Gao, Hao Wu, Tianyi Li, Runhua Liu, Yu Sun, Jiaqi Wang, Linlin Fang, Chenning Zhang, Yikun Sun

## Abstract

Jing-Fu-Kang granules (JFKG) is a famous Chinese patent medicine for the treatment of cervical spondylosis around the China, whereas the active substance and mechanism are not completely investigated clearly. In the current study, a rapid separation and identification method using UPLC-QE-Orbitrap-MS was established, 97 chemical constituents from JFKG were identified, and 16 prototype components from plasma samples after administration of JFKG were observed within 16 min. The structures of typical compounds were preliminarily speculated by comparing the retention time and fragmentation pattern. Furthermore, multiple databases were used to integrate the compound targets of JFKG, and the disease targets related to cervical spondylosis. After the intersection of the two sets of targets, a protein-protein interaction (PPI) network and a TCM-component-target-pathway-disease network were established, then using the DAVID database to perform gene ontology analysis and Kyoto Encyclopedia of Genes and Genomes analysis on the common targets to find related pathways. Finally, a total of 531 common targets and 136 pathways were found to participate in the mechanism. Our findings will help to further confirm the mechanism of JFKG for relieving cervical spondylosis, which will improve the scientific rationality of JFKG in clinical use, and can also assist in guiding doctors.

## 1. Introduction

Cervical spondylosis (CS) is a common disease that affects cervical vertebrae at many levels, including intervertebral disc degeneration, vertebral osteophytes, facet and pedicle hypertrophy, and ligament and segmental instability [1]. As far as the symptoms of CS are concerned, it is equivalent to the category of “nape arthralgia” and “stiff neck” in traditional Chinese medicine, which is mainly characterized by neck pain, clonus, shoulder, arm numbness and so on [2]. Epidemiological investigation showed that the incidence of CS is increasing year by year and getting younger, which seriously affects the patients’ quality of life [3]. Most CS can be treated by non-operative treatment, and only a few patients with CS who are ineffective by long-term systematic non-operative treatment or without contraindications need surgical treatment [4]. Therefore, drug therapy is the mainstream method for the treatment of CS.

Jing-Fu-Kang granules (JFKG) is developed and produced by Chengde Jing fu kang Pharmaceutical Group (Hebei, China), including 21 kinds of traditional Chinese medicines, such as Puerariae lobatae Radix, Gentianae macrophyllae Radix, Atractylodes Rhizome, Salviae miltiorrhizae Radix et Rhizome, Paeoniae Radix Alba, Carthami Flos, Astragali Radix, Phellodendri chinensis Cortex, etc. JFKG own the effect of promoting blood circulation, dredging collaterals, dispelling wind and relieving pain. It can not only improve the elasticity of vertebral artery, relieve vasospasm, improve cerebral blood circulation, but also relieve muscle and ligament spasm and reduce the compression of blood vessels and nerves. At the same time, it can improve other symptoms of CS, such as neck stiffness, limited movement, neck and shoulder pain, numbness and weakness of limbs, and the total effective rate can reach more than 90%. Without a doubt, JFKG is the first choice for the treatment of CS, which can remove the etiology from the root and not easily to repeat the disease of using with a long time and improve the quality of life of patients [5–7].

Due to the complex composition and diversified mechanism of traditional Chinese medicine, the progress of the research on the material basis of traditional Chinese medicine is difficult, which seriously affects the modernization of traditional Chinese medicine [8–9]. As for the material basis of traditional Chinese medicine formulas, we should understand the composition of the formulas firstly, then determine the components in the body to provide information for the study of the whole medicine formulas and this will also become one way for the research of traditional Chinese medicine formulas [10]. In fact, the effective material basis of traditional Chinese medicine and traditional Chinese medicine formulas can be further determined by the chemical composition in the plasma after oral administration and this method will be faster and more accurate than the traditional analytical method [11]. However, At present, there are few studies on JFKG, and no scientist has conducted a comprehensive study on the chemical components of JFKG, but only several compounds were evaluated in JFKG. For example, Yujie Yang [12] showed that the volatile oil of JFKG has significant anti-inflammatory and analgesic effects, and its mechanism may be related to the decrease of the levels of IL-1, TNF-α and PGE2. Therefore, in this experiment, the chemical components from JFKG were comprehensively analyzed, and the compounds absorbed into the blood after administration JFKG was explored for the first time.

Recent years, ultra-high performance liquid chromatography-tandem mass spectrometry (UPLC-MS) has been increasingly used in the rapid analysis of complex traditional Chinese medicine chemical systems because of its high resolution, high sensitivity, and high selectivity [13–17]. What’s more, with the rapid development of bioinformatics, network pharmacology has become a powerful tool for exploring traditional Chinese medicine formulas, which is a perfect method to investigate drug mechanism, determine new targets and expand new indications at a systematic level. It can fully reflect the action mechanism of drugs on the disease network, providing support for drug discovery of complex diseases and natural products [18–19]. Overview, in this article, UPLC-QE-Orbitrap-MS skill and network pharmacology method were used to for the first time, combined with relevant literatures and chemical mass spectrum information, JFKG effective extraction constituents and plasma samples after administration of JFKG effective extraction were quickly analyzed and identified, and GO, KEGG analysis was carried out. It provides a research basis for clarifying the material basis of pharmacodynamics, and elucidating the mechanism of pharmacological action in vivo, which will help to improve the level of quality control, and ensuring the safety and effectiveness of clinical drug use.

## 2. Materials and methods

### 2.1. Materials and Reagents

JFKG (Lot number: 20111005); Acetonitrile (MS grade); formic acid (MS grade); methanol (MS grade) [all three reagents were purchased from Thermo Fisher Scientific (China) Co., Ltd.]; ultrapure water (ultra-purified by Milli-Q Advantage A10 Water system).

### 2.2. Animals

6 male Sprague-Dawley rats (320-345g) were purchased from the Si pei fu (Beijing, China) Biotechnology Co., Ltd. There were three animals in both the experimental and blank groups. Rats were housed in an animal room (25 °C, 60±5% humidity, 12-h dark-light cycle) with free access to water and normal diet ad libitum. All the animals were bred under the above conditions for 4-day acclimation and fasted overnight before drug administration. The animal experiment protocol was approved by the Animal Care and Use Committee of Beijing University of Chinese Medicine. The JFKG sample was dissolved in distilled water. Rats were randomized by body weight and orally administered under the dose of 6.84 g/kg. At the same time, the blank group was orally administered with the same dose of distilled water. 3 rats in the experimental group were given the suspension prepared by intragastric administration at a dose of 6.84 g/kg, and 3 rats in the blank group were intragastrical drunk twice a day for 3 days. After the last administrations, a 0.2 mL volume of blood was collected in a heparinized microcentrifuge tube from tail vein at the following time points: 15 min, 30 min, 1 h, 2 h, 4 h, 6 h, and then centrifuged at 4000 rpm for 10 min; the supernatant was frozen at −80 °C before analysis.

### 2.3. Preparation of JFKG extract

Accurately weighed granules (1.01 g) was suspended in 10 mL 80% (v/v) methanol-water and ultrasonically extracted for 30 min and then cooled to room temperature. After centrifugation at 10000 rpm for 5 min, to get 50 μL supernatant, diluting 20 times with 80% (v/v) methanol-water, centrifugation 13000 rpm for 15 min to obtain the sample solution.

### 2.4. Preparation of plasma samples

For analysis, plasma samples frozen at −80°Cwere thawed at room temperature. Different time points plasma samples were mixed in equal amounts. 360 μL of methanol solution was added to 120 μL of the mixed plasma sample, vortexed for 3 min, sonicated in an ice-filled ultrasonic water bath for 10 minutes, and centrifuged at 4000 rpm for 10 min. The supernatant was taken and blown dry at 40 °C under nitrogen. The residue was dissolved by 120 μL of methanol, vortexed for 30 s and centrifuged at 12000rpm for 5 min, and the injection volume was 5 μL.

### 2.5. Main instruments

Thermo Scientific Q Exactive (Thermo Scientific); Xcalibur (Thermo Scientific); Vanquish Duo UHPLC System for Dual LC Workflows (Thermo Scientific); Waters BEH UPLC C18 column( 1.7 μm, 2.1×150 mm, Milford, MA, USA );CPA225D Electronic balance (SARTORIUS); KQ5200DA Ultrasonic Cleaners (Kunshan Shumei, China); Pico 17 High speed centrifuge( Thermo Scientific ); Nitrogen evaporator N-EVAP 116 (Organomation).

### 2.6. UPLC-QE-Orbitrap-MS Conditions

Waters ACQUITY UPLC BEH C18 column (1.7 um, 2.1×150 mm, Milford, MA, USA), column temperature 40 °C, injection volume 5 μL, flow rate 0.3 ml/min, mobile phase: 0.1% formic acid aqueous solution (A) and acetonitrile (B). The eluting program was: 5-5%B for 0-2 min, 5-98%B for 2-16min.

The ion source is electron spray ionization (ESI); MS was operated in negative/positive mode; scan mode was Full scan/ddMS2, positive and negative ion alternating scanning, and the scanning mode is 100-1300 Da, capillary temperature 350 °C. The spray voltage in positive mode is 3200 V, the spray voltage in negative mode is 3800 V, the sheath gas is 35 arb, and the auxiliary gas is 15 arb. Three collision energies of low, medium and high are used for MS2. The positive ion mode is 20 V, 40 V, 60 V, and the negative ion mode is 30 V, 50 V, 70 V. Resolution of primary mass spectrometry Full Scan 70000 FWHM (Full Width at Half Maximum), secondary mass spectrometry resolution: MS/MS 17500 FWHM.

### 2.7. Compound Identification

According to the chromatographic retention time, relative molecular mass, fragment ion information, relevant literature data, and MS fragmentation ion, JFKG effective extraction constituents and plasma samples after administration of JFKG were identified.

### 2.8. Network Pharmacology Construction

#### 2.8.1. Active ingredient screening

The identified compounds were screened on the condition of Intensity>10^7^ in the ion flow diagram. In order to avoid the screening of individual active components with high recognition, the active components were obtained by consulting the relevant literature of Chinese Pharmacopoeia (2015 edition) and CNKI at the same time. The target is predicted by Swiss Target Prediction database (http://www.swisstargetprediction.ch/).

#### 2.8.2. Prediction of disease targets of cervical spondylosis

Enter the key word “cervical spondylosis/ Cervical spondylopathy/ cervical syndrome” into the human OMIM database (https://omim.org/) and GeneGards database (https://www.genecards.org/) to obtain the disease target.

#### 2.8.3. Acquisition of common targets for component diseases and construction of protein-protein interaction network

The target of JFKG and the target of disease were intersected by Venn map of WeChat website (http://www.bioinformatics.com.cn/), and the common target of sub-disease was obtained. The common target points were imported into STRING database (https://string-db.org/cgi/input.pl) to draw Protein-Protein Interaction (PPI), and Cytoscape3.8.2 software was used to visualize the structure of protein network and analyze its topological characteristics. Taking the median of Degree Centrality (DC), Betweenness Centrality (BC) and Closeness Centrality (CC) as screening conditions, the core targets of component therapy were obtained twice.

#### 2.8.4. GO analysis and KEGG enrichment analysis

we enriched the hub genes by Gene Ontology (GO) analysis and Kyoto Encyclopedia of Genes and Genomes (KEGG) pathway enrichment analysis in the Database for Annotation, Visualization, and Integrated Discovery (https://david.ncifcrf.gov), and the results were visualized with the bioinformatics online tool (http://www.bioinformatics.com.cn/).

#### 2.8.5. Network construction

The effective components, core targets and action pathways of JFKG were introduced into Cytoscape3.8.2 software to establish a “TCM-component-target-pathway-disease” network.

## 3. Results

### 3.1. UPLC-QE-Orbitrap-MS analysis of JFKG constituents

#### 3.1.1. Chemical composition analysis

Considering the complex chemical components, the components of JFKG were fully scanned by positive and negative ionization modes. The retention time and mass spectrometry information of compounds from JFKG were obtained by QE detection. Based on the obtained accurate mass spectrometry information such as fragment ions, reference substance and the reference literatures, 97 chemical constituents were identified in JFKG. The results are shown in Tab 1, the Base peak intensity chromatogram (BPI) is shown in Fig 1.

**Fig 1.**
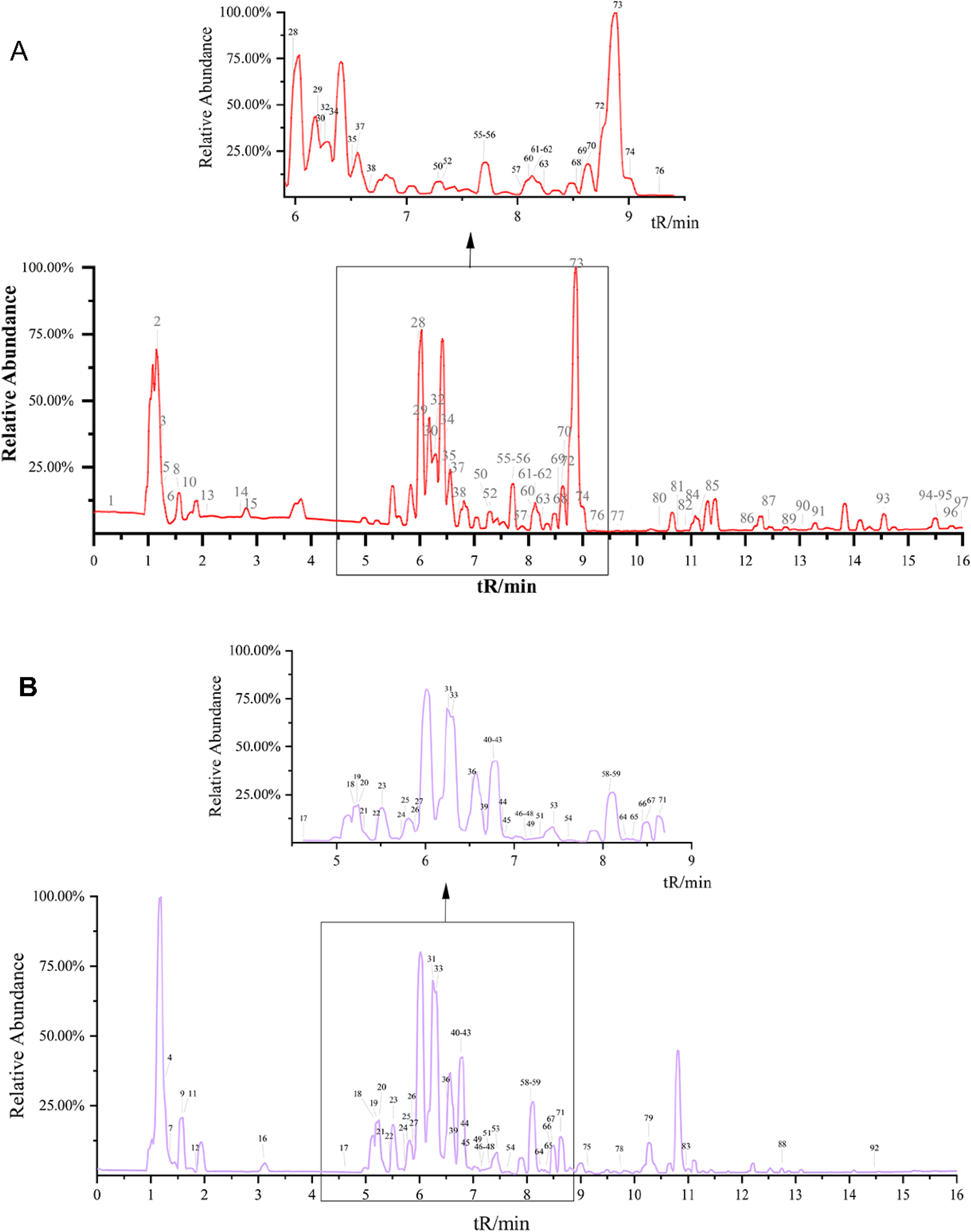
Base peak ion flow diagram of Jing-Fu-Kang granules under positive ion ( A ) and negative ion ( B ) from UPLC-QE-Orbitrap-MS.

**Table 1.**
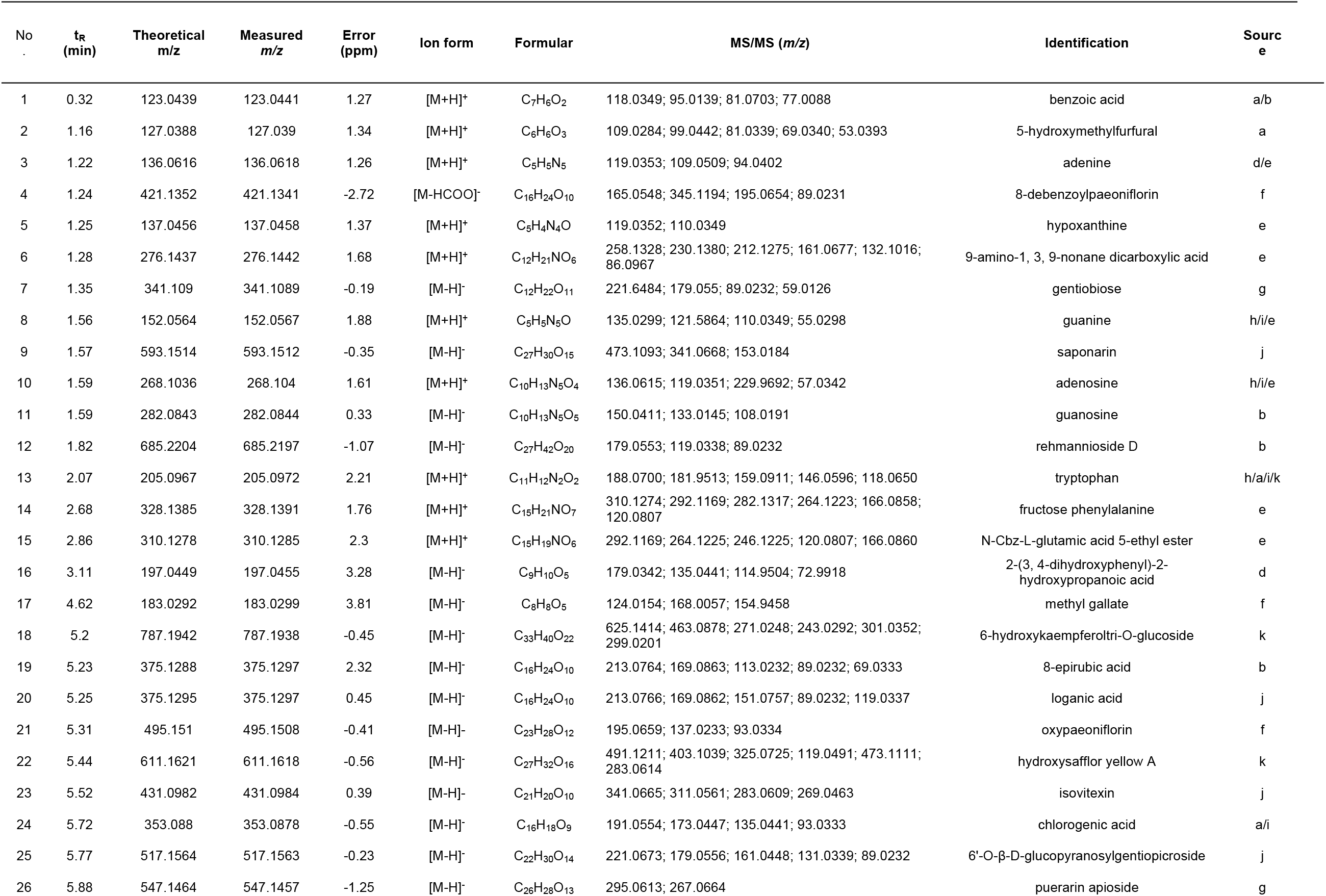
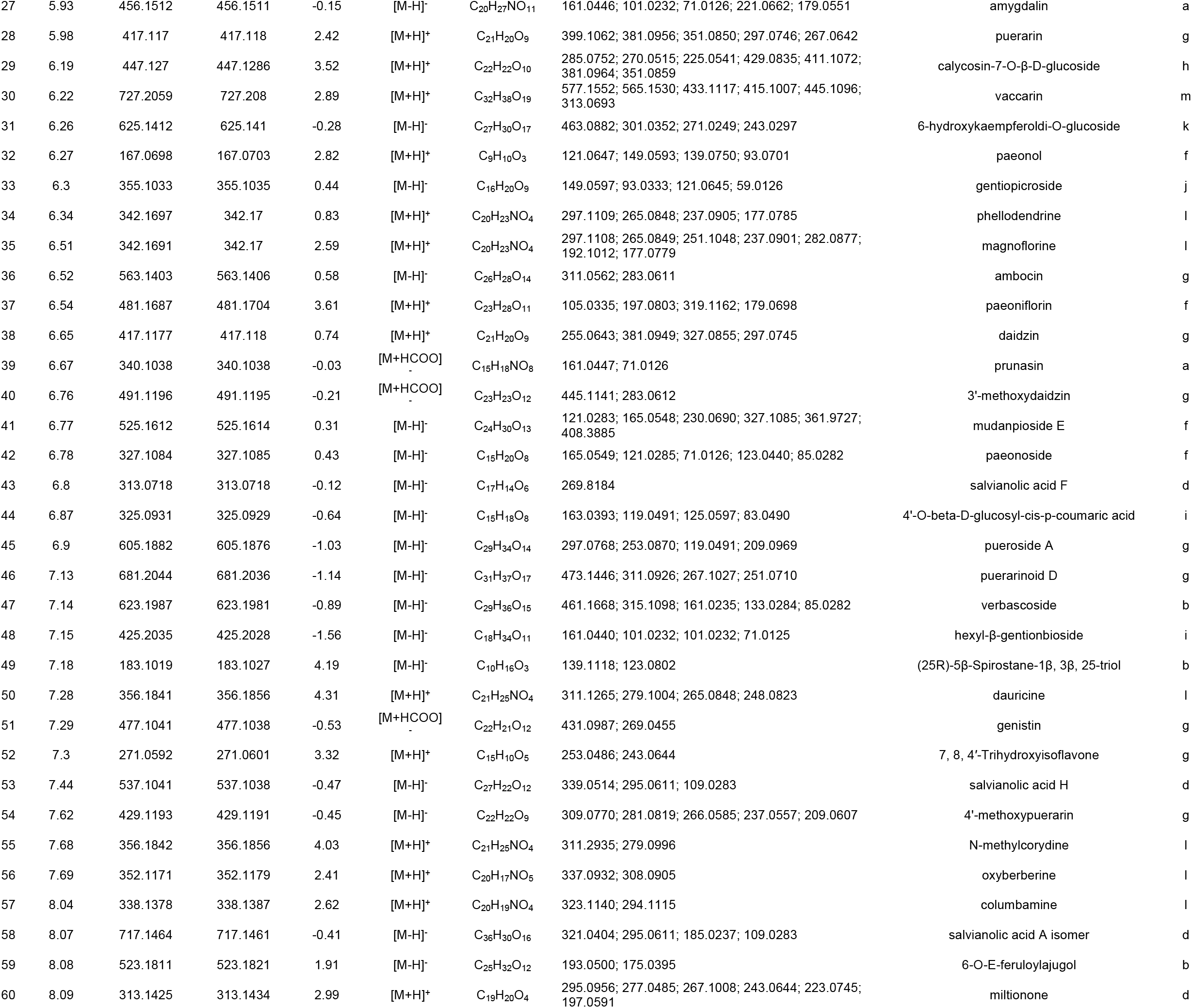
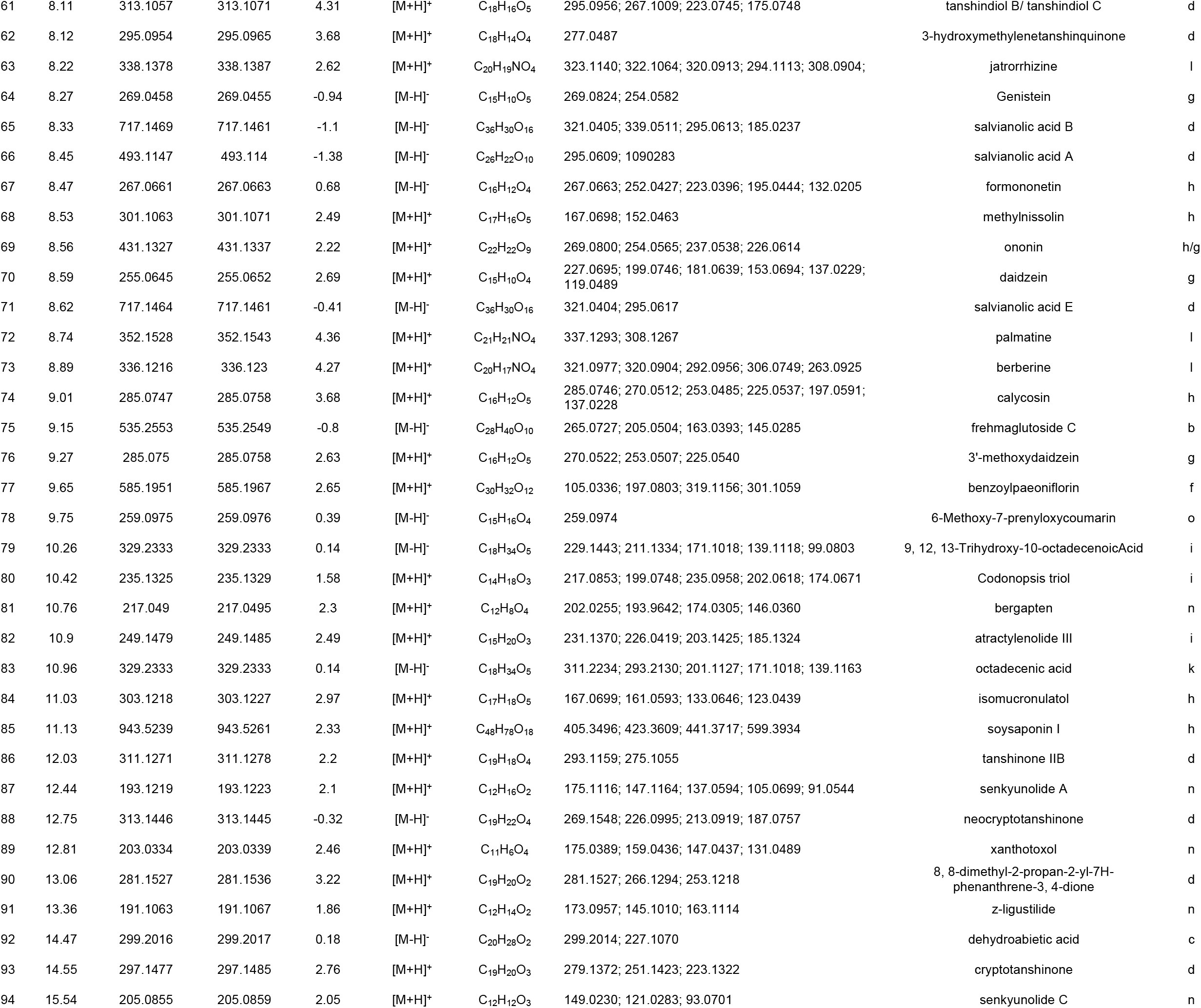
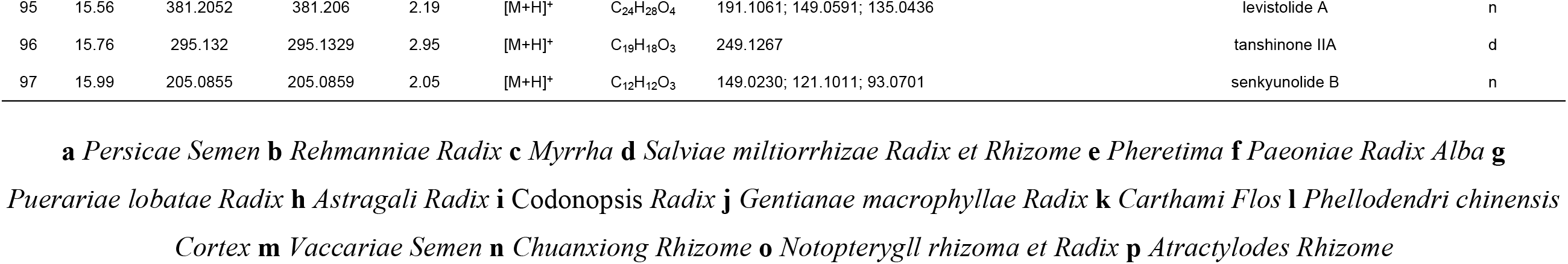
The identified JFKG effective extraction constituents by UPLC-QE-Orbitrap-MS.

#### 3.1.2. Identification of known components

The retention time of peak 73 is 8.89 min, and the excimer ion peak *m/z* 336.1216 can be observed. At the same time, the characteristic fragments of *m/z* 321.0977 [M+H-CH_3_]^+^, *m/z* 320.0904 [M+H-CH_3_-H]^+^, *m/z* 306.0749 [M+H-CH_2_O]^+^, *m/z* 292.0956 [M+H-CH_3_-H-CO]^+^ can be observed. According to the *m/z* in the positive ion mode and the mass of the characteristic fragments, combined with the related literature [17–18], the compound can be judged to be berberine from the *Cortex Phellodendri* (Fig 2A).

**Fig 2.**
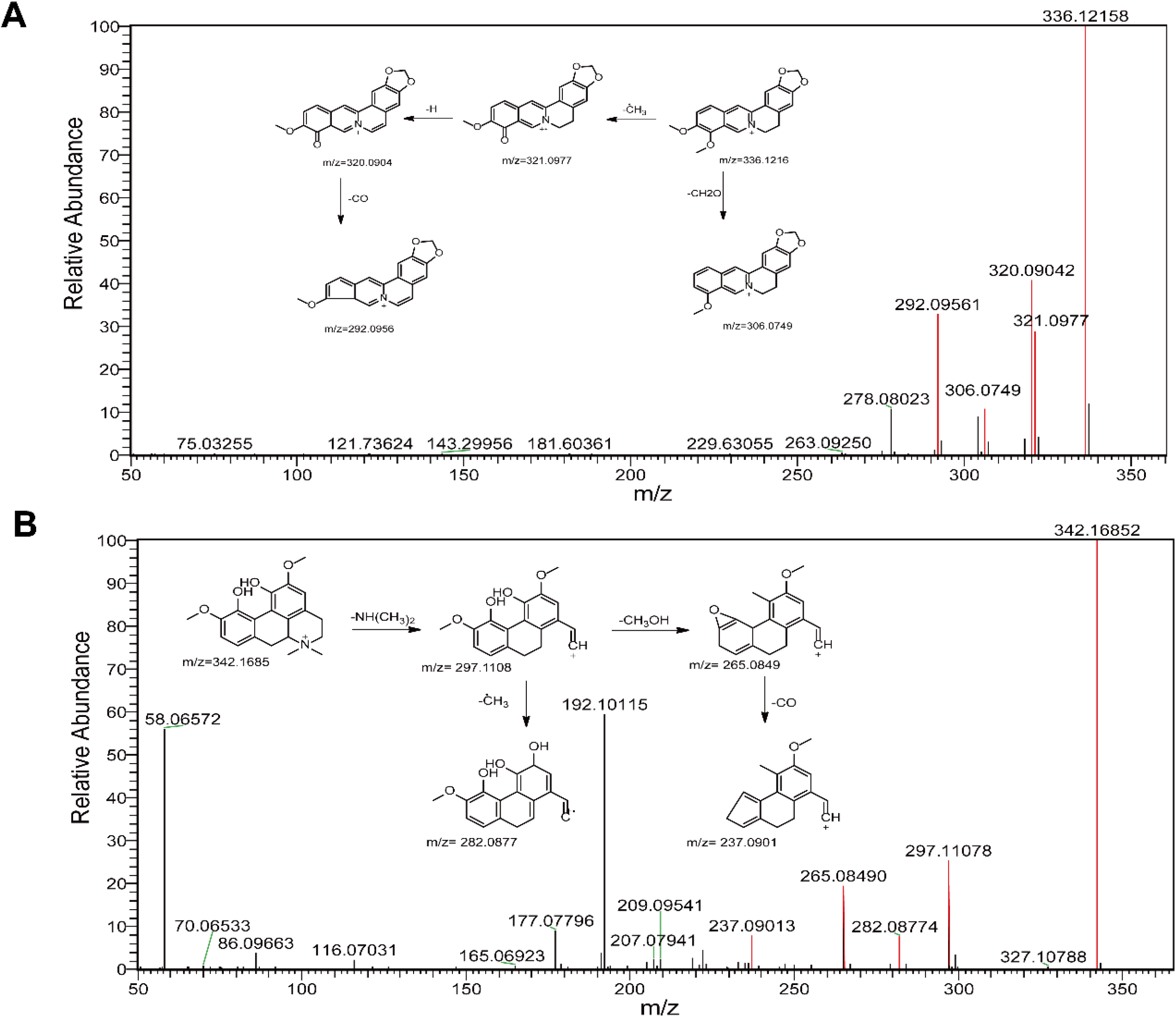
The MS/MS spectrum of berberine (A), magnoflorine (B) in positive ion mode and probable fragmentation pathway.

The retention time of peak 37 is 6.51 min, and the excimer ion peak *m/z* 342.1685 [M+H]^+^ can be observed. At the same time, the characteristic fragments *m/z* 297.1108 [M+H-C_2_H_7_N]^+^, *m/z* 282.0877 [M+H-C_2_H_7_N-CH_3_]^+^, *m/z* 265.0849 [M+H-C_2_H_7_N-CH_3_-OH]^+^, *m/z* 237.0901 [M+H-C_2_H_7_N-CH_3_-OH-CO]^+^ also can be observed. According to the *m/z* in the positive ion mode and the quality of the characteristic fragments, combined with the related literatures [17–18], the compound can be judged to be magnoflorine from the *Cortex Phellodendri* (Fig 2B).

The retention time of peak 28 is 5.98 min, and the excimer ion peak *m/z* 417.1166 [M+H]^+^ can be observed. At the same time, the characteristic fragments 399.1062 [M+H-H_2_O]^+^, 381.0956 [M+H-2H_2_O]^+^, 351.0850 [M+H-2H_2_O-CH_2_O]^+^, 297.0746 [M+H-H_2_O-C_4_H_6_O_3_]^+^ also can be observed. According to the *m/z* in the positive ion mode and the quality of the characteristic fragments, combined with the related literatures [16–17,19], the compound can be judged to be puerarin from the *Puerariae lobatae Radix.*

The retention time of peak 21 is 5.31 min, the first level produces [M-H]^−^ peak of *m/z* 495.1521, and the second stage cleavage produces fragments of 195.0659 [M-H-C6H10O5-C7H6O3]^−^, 137.0233 [M-H-C6H10O5-C10H12O4]^−^, 93.0334 [M-H-C6H10O5-C10H12O4-CO2]^−^. According to the *m/z* in the negative ion mode and the quality of the characteristic fragments, combined with the related literature [20], the compound can be determined to be oxypaeoniflorin from the *Paeoniae Radix Alba.*

The retention time of peak 20 is 5.25 min, the first level produces [M-H]-peak of *m/z* 375.1301, and the second stage cleavage produces fragments of 213.0766 [M-H-C6H11O5]^−^, 169.0862 [M-H-C6H11O5-CO2]^−^, 119.0337 [M-H-C6H11O5-CO2-CH2-2H2O]^−^. According to the *m/z* in the negative ion mode and the quality of the characteristic fragments, combined with the related literature [21], the compound can be determined to be loganic acid (Fig 3A) from the *Gentianae macrophyllae Radix.*

**Fig 3.**
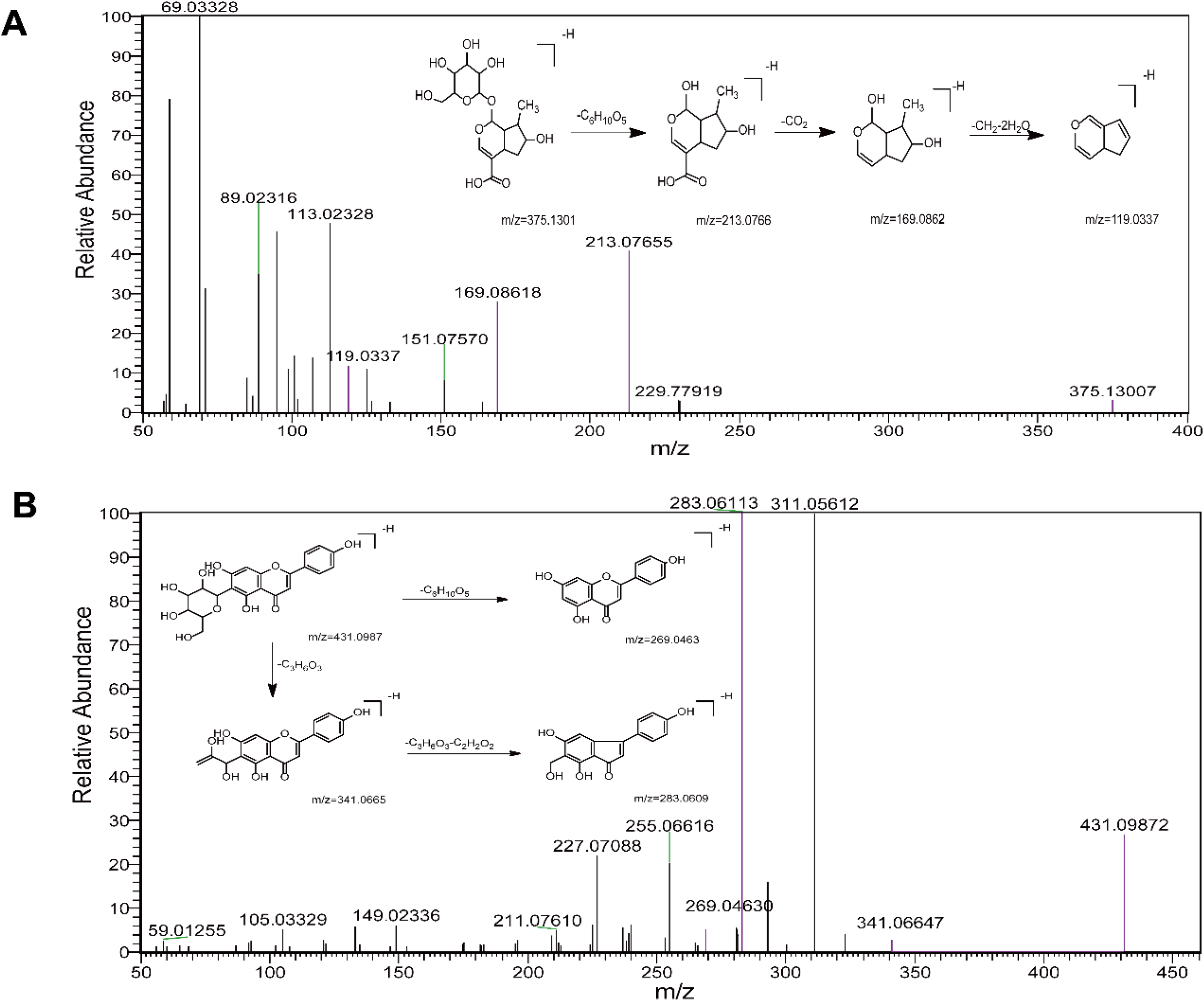
The MS/MS spectrum of loganic acid (A), isovitexin (B) in negative ion mode and probable fragmentation pathway.

The retention time of peak 23 is 5.52 min, the first level produces [M-H]-peak of *m/z* 431.0987, and the second stage cleavage produces fragments of 341.0665 [M-H-C3H6O3]^−^, 283.0609 [M-H-C3H6O3-C2H2O2]^−^, 269.0463 [M-H-C6H10O5]^−^. According to the *m/z* in the negative ion mode and the quality of the characteristic fragments, combined with the related literature [21], the compound can be determined to be isovitexin (Fig 3B) from the *Gentianae macrophyllae Radix.*

The retention time of peak 93 is 12.44 min, and the excimer ion peak *m/z* 193.1218 [M+H]^+^ can be observed. At the same time, the characteristic fragments 175.1116 [M+H-H2O]^+^, 147.1164 [M+H-H2O-CO]^+^, 137.0594 [M+H-C4H8]^+^, 105.0699 [M+H-H2O-CO-C3H6]^+^ also can be observed. According to the *m/z* in the positive ion mode and the quality of the characteristic fragments, combined with the related literature [22], the compound can be judged to be senkyunolide A from the *Chuanxiong Rhizome*.

The retention time of peak 47 is 7.14 min, the first level produces [M-H]^−^ peak of *m/z* 623.1986, and the second stage cleavage produces fragments of 461.1668 [M-H-C9H6O3]^−^, 161.0235 [M-H-C9H6O3-C14H20O7]^−^. According to the *m/z* in the negative ion mode and the quality of the characteristic fragments, combined with the related literature [19], the compound can be determined to be verbascoside from the *Rehmanniae Radix.*

The retention time of peak 22 is 5.43 min, the first level produces [M-H]^−^ peak of *m/z* 611.1615, and the second stage cleavage produces fragments of 491.1211 [M-H-C4H8O4]^−^, 473.1112 [M-H-C4H8O4-H2O]^−^, 403.1039 [M-H-C4H8O4-C3H4O3]^−^, 325.0725 [M-H-C4H8O4-H2O-C4H4O6]^−^, 283.0614 [M-H-C4H8O4-H2O-C4H4O6-C2H2O]^−^, 119.0491 [M-H-C19H24O15]^−^. According to the *m/z* in the negative ion mode and the quality of the characteristic fragments, combined with the related literatures [18, 23] the compound can be determined to be hydroxysafflor yellow A from the *Carthami Flos.*

### 3.2. UPLC-QE-Orbitrap-MS analysis of the components absorbed into the blood after administration of JFKG extract

Components absorbed into the blood are always considered to be active substances with potential pharmacological effects. Therefore, the plasma samples of rats after oral administration of JFKG were analyzed by UPLC-QE-Orbitrap-MS. By comparing the retention time, fragment ions of plasma samples after administration of JFKG extract and blank plasma, and the previous identification of the chemical composition of JFKG extract, a total of 16 prototype components were identified with an error of 5ppm. The BPI of rat blank plasma and rat plasma samples after oral administration JFKG are shown in fig 4. The mass spectrometric data of 16 prototype components are summarized in Table 2. Fig 4 show the BPI of rat mixed blank plasma and rat mixed plasma samples after oral administration, and the classification of chemical components and blood plasma are shown in Fig 5.

**Table 2.** The identified chemical constituents from plasma samples after administration of JFKG by UPLC-QE-Orbitrap-MS.

| No. | tR<br>(min) | Theoretical<br>(*m/z*) | Measured<br>(*m/z*) | Error<br>(ppm) | Ion form | Formular | MS/MS(*m/z*) | Identification | Source |
| --- | --- | --- | --- | --- | --- | --- | --- | --- | --- |
| 1 | 1.63 | 268.1036 | 268.1033 | -1.12 | [M+H]<sup>+</sup> | C<sub>10</sub>H<sub>13</sub>N<sub>5</sub>O<sub>4</sub> | 268.1038; 136.0615; 57.0341 | adenosine | h/i/e |
| 2 | 2.63 | 328.1385 | 328.1381 | -1.22 | [M+H]<sup>+</sup> | C<sub>15</sub>H<sub>21</sub>NO<sub>7</sub> | 310.1275; 292.1169; 264.1221; 120.0806; 166.0857 | fructose phenylalanine | e |
| 3 | 2.77 | 310.1278 | 310.1277 | -0.32 | [M+H]<sup>+</sup> | C<sub>15</sub>H<sub>19</sub>NO<sub>6</sub> | 292.1164; 264.1220; 246.1115; 166.0858; 120.0805 | N-Cbz-L-glutamic acid 5-ethyl ester | e |
| 4 | 5.89 | 417.1170 | 417.1176 | 1.44 | [M+H]<sup>+</sup> | C<sub>21</sub>H<sub>20</sub>O<sub>9</sub> | 381.0963; 321.0749; 297.0749; 267.0645 | puerarin | g |
| 5 | 5.96 | 547.1464 | 547.1456 | -1.46 | [M-H]<sup>-</sup> | C<sub>26</sub>H<sub>28</sub>O<sub>13</sub> | 267.0660; 295.0608 | puerarin apioside | g |
| 6 | 6.30 | 167.0698 | 167.0703 | 2.99 | [M+H]<sup>+</sup> | C<sub>9</sub>H<sub>10</sub>O<sub>3</sub> | 121.4165; 149.0964; 139.0753; 93.0701 | paeonol | f |
| 7 | 6.35 | 342.1697 | 342.1697 | 0 | [M+H]<sup>+</sup> | C<sub>20</sub>H<sub>23</sub>NO<sub>4</sub> | 342.1691; 297.1114; 265.0853; | phellodendrine | l |
| 8 | 8.00 | 338.1378 | 338.1381 | 0.89 | [M+H]<sup>+</sup> | C<sub>20</sub>H<sub>19</sub>NO<sub>4</sub> | 338.1378; 323.1145; 294.1117; | columbamine | l |
| 9 | 8.04 | 313.1057 | 313.1065 | 2.56 | [M+H]<sup>+</sup> | C<sub>18</sub>H<sub>16</sub>O<sub>5</sub> | 313.1550; 295.0955; 267.1006; 249.0907; 107.0493 | tanshindiol B/ tanshindiol C | d |
| 10 | 8.56 | 255.0645 | 255.0647 | 0.78 | [M+H]<sup>+</sup> | C<sub>15</sub>H<sub>10</sub>O<sub>4</sub> | 255.0646; 227.0698; 199.0751; 181.0647; 157.0643; 137.0231 | daidzein | g |
| 11 | 8.69 | 352.1528 | 352.1535 | 1.99 | [M+H]<sup>+</sup> | C<sub>21</sub>H<sub>21</sub>NO<sub>4</sub> | 352.1535; 336.1223; 308.1274; 321.0988; 320.1269 | palmatine | l |
| 12 | 8.98 | 285.0747 | 285.0754 | 2.46 | [M+H]<sup>+</sup> | C<sub>16</sub>H<sub>12</sub>O<sub>5</sub> | 285.1086; 212.0803 | calycosin | h |
| 13 | 9.66 | 259.0975 | 259.0973 | -0.77 | [M-H]<sup>-</sup> | C<sub>15</sub>H<sub>16</sub>O<sub>4</sub> | 259.0975; 215.1072 | 6-Methoxy-7-prenyloxycoumarin | o |
| 14 | 10.20 | 329.2333 | 329.2332 | -0.3 | [M-H]<sup>-</sup> | C<sub>18</sub>H<sub>34</sub>O<sub>5</sub> | 329.2334; 293.2125; 261.1133; 211.1334; 171.1018; 139.1118; 99.0803 | 9;12;13-Trihydroxy-10-octadecenoicAcid | i |
| 15 | 11.04 | 329.2333 | 329.2331 | -0.61 | [M-H]<sup>-</sup> | C<sub>18</sub>H<sub>34</sub>O<sub>5</sub> | 329.2337; 311.2234; 293.2125; 199.1333; 171.1019 | octadecenic acid | k |
| 16 | 12.75 | 313.1446 | 313.1448 | 0.64 | [M-H]<sup>-</sup> | C<sub>19</sub>H<sub>22</sub>O<sub>4</sub> | 313.2384; 213.0914; 201.1126; 187.0748; | neocryptotanshinone | d |

**Fig 4.**
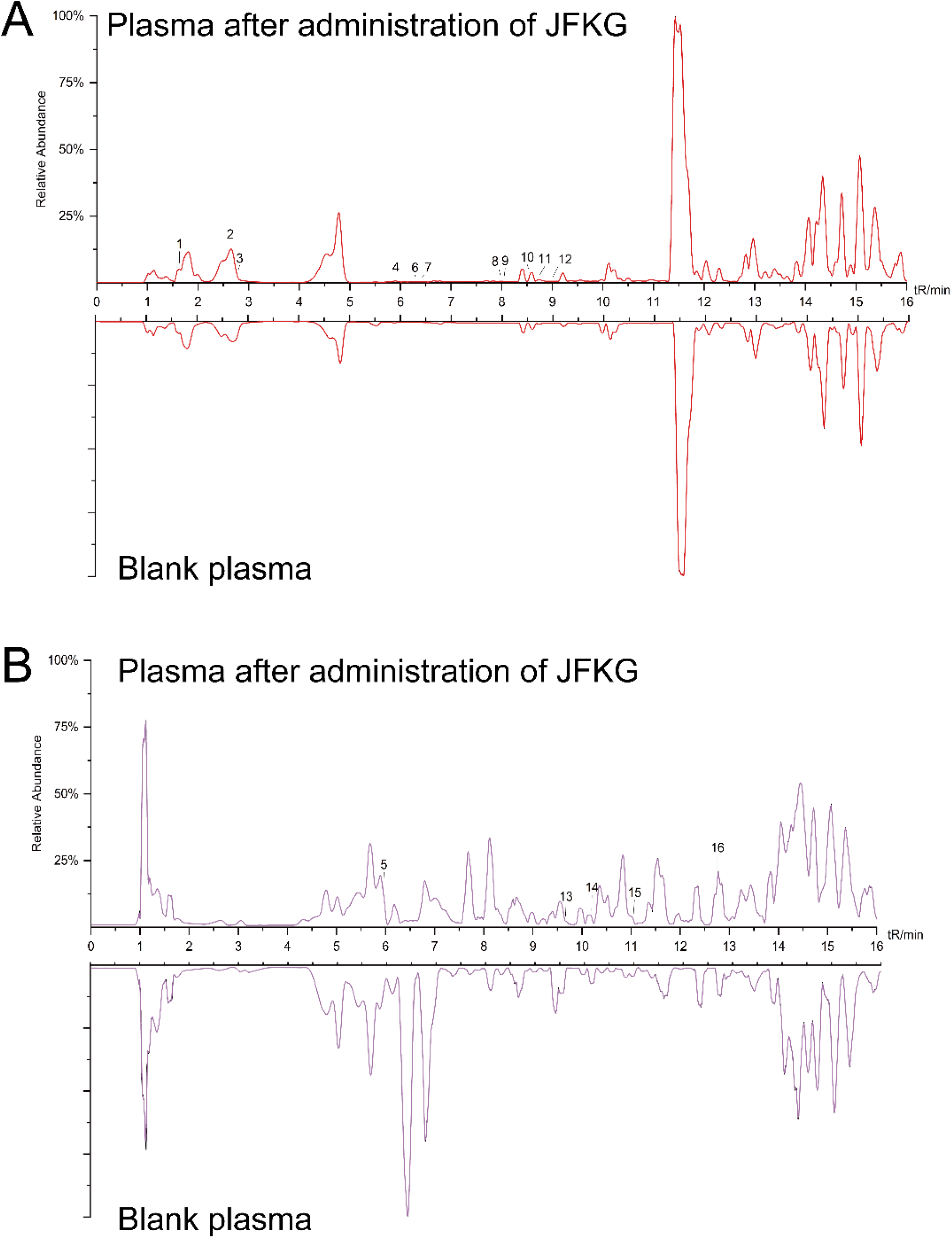
BPI of plasma samples after administration of JFKG and blank plasma positive ion (A) and negative ion (B) by UPLC-QE-Orbitrap-MS.

**Fig 5.**
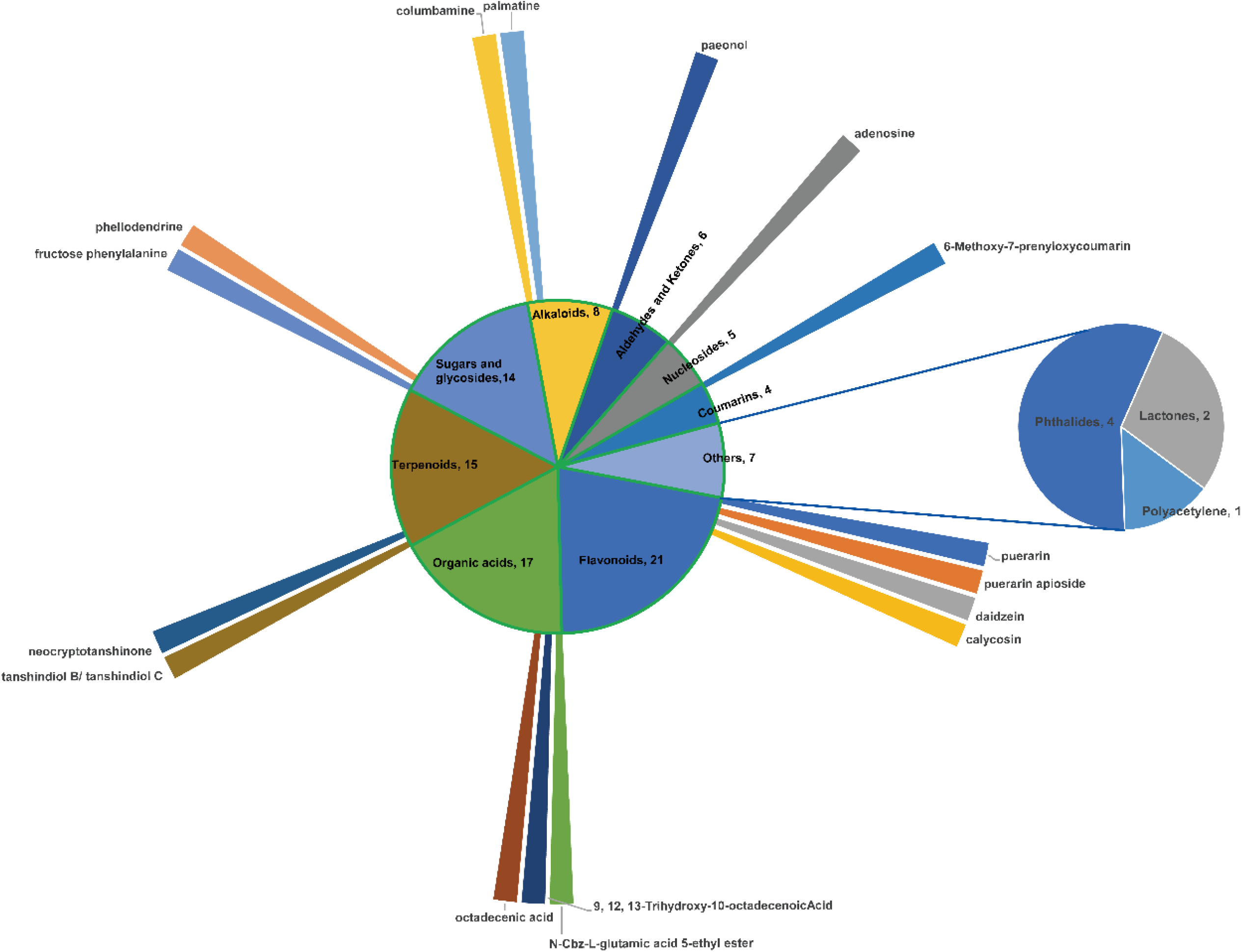
classification of the identified chemical components from JFKG extract and blood components.

### 3.3. Network pharmacology

#### 3.3.1. Screening and prediction of JFKG targets

The identified compounds were screened under the condition of intensity > 10^7^ in the ion current diagram; and 62 compounds were selected as target chemical constituents by consulting the relevant literature of Chinese Pharmacopoeia (2015 edition) and CNKI. Using the Swiss Target Prediction website to predict targets; a total of 2818 targets were obtained.

#### 3.3.2. Target prediction of cervical spondylosis

Using the keyword “cervical spondylosis/ Cervical spondylopathy/ cervical syndrome” to search for disease targets in OMIM and GeneGards databases; screening and merging to remove duplicates; a total of 6923 disease targets were obtained.

#### 3.3.3. PPI construction and topological analysis

The Venn map is made through WeChat website(http://www.bioinformatics.com.cn/); and 531 common targets are obtained (Fig 6).

**Fig 6.**
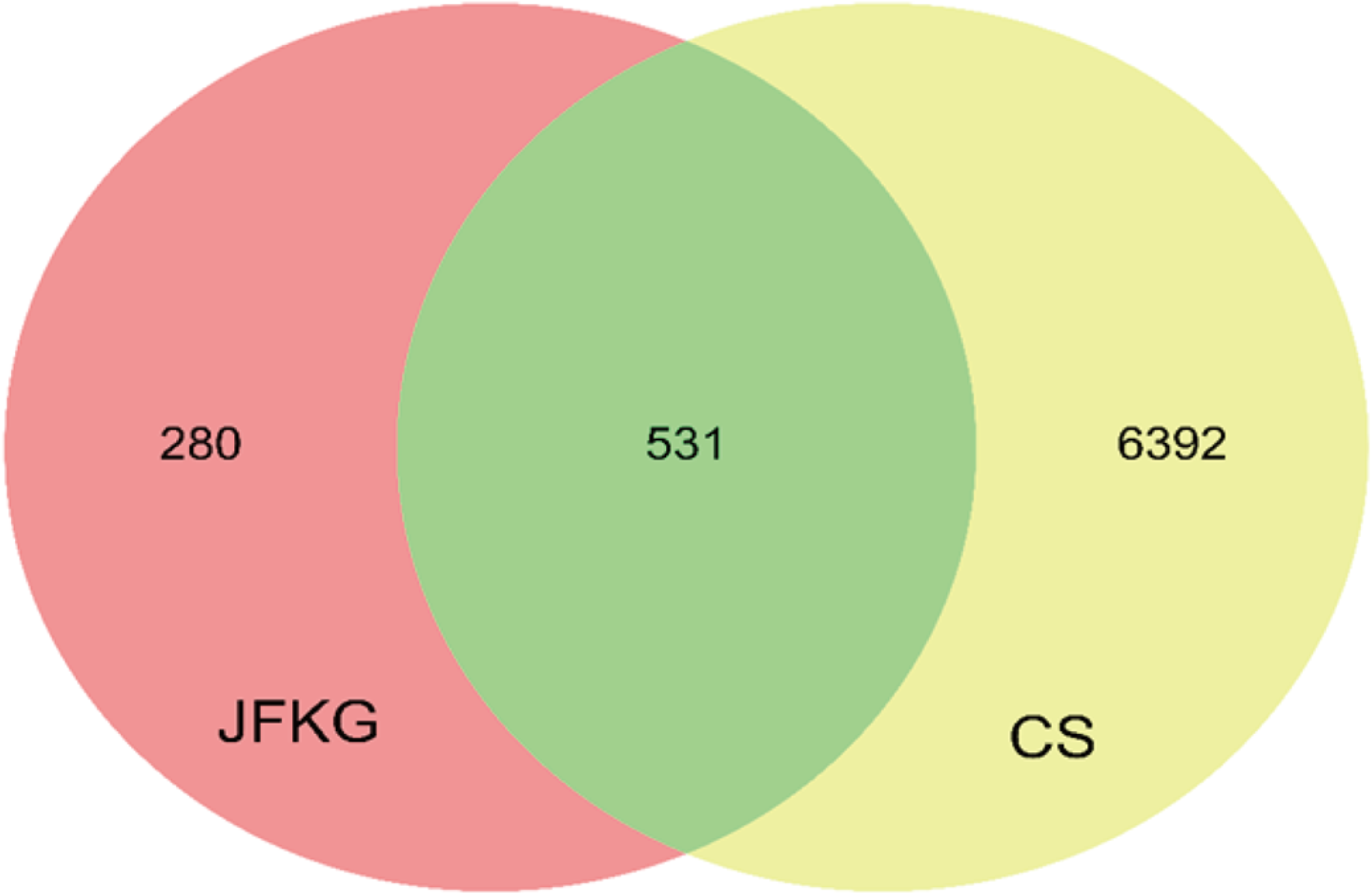
The Venn diagram of common targets from the components of extraction of JFKG and disease.

The String website is a network of functional protein associations that can predict protein-protein interactions. After all the targets were imported into the website; the PPI network map was obtained and saved in tsv format. In this study; medium confidence > 0.9 was set; edge free proteins were excluded; and the reliability of the research data was ensured. The tsv file was exported for backup. The topological characteristics of the protein network structure were analyzed by Cytoscape3.8.2 software; and 13 core targets for the treatment of CS were screened (Fig 7 and Table 3).

**Table 3.**
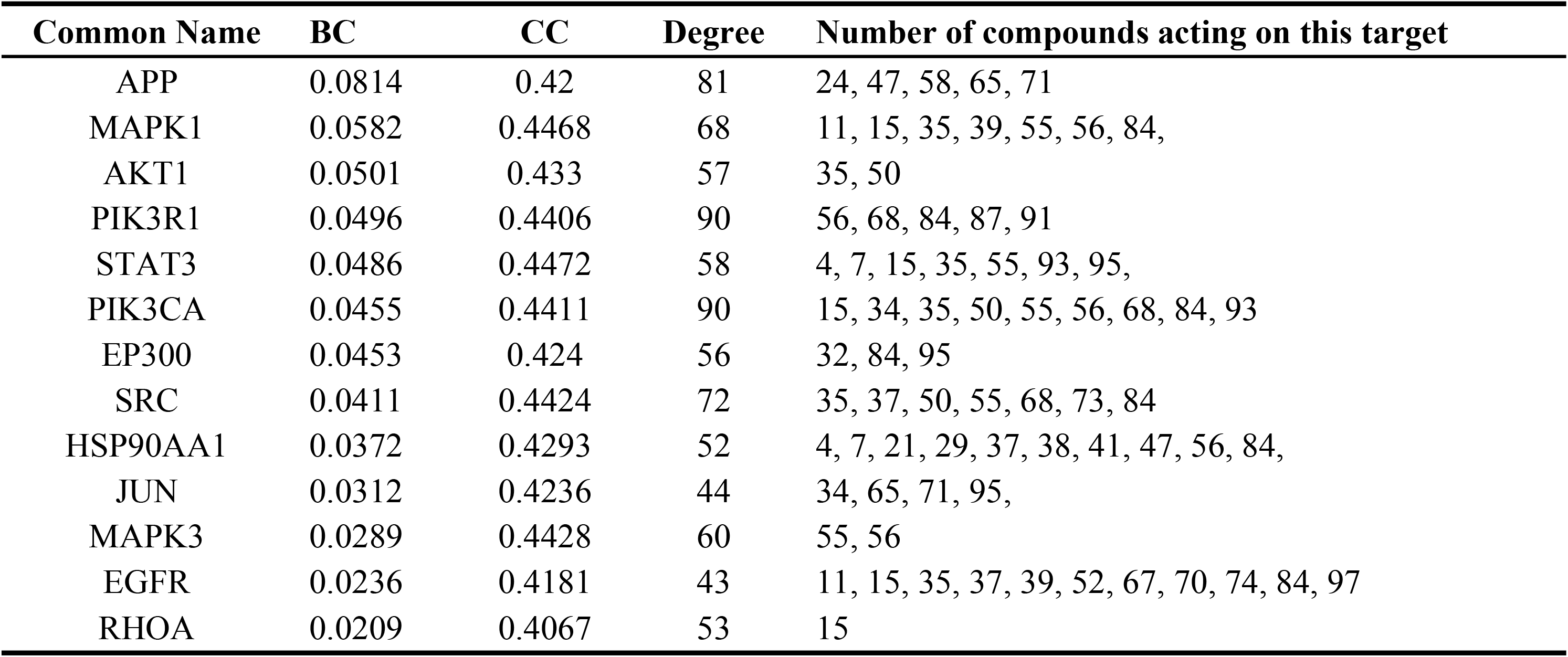
Core targets of information.

**Fig 7.**
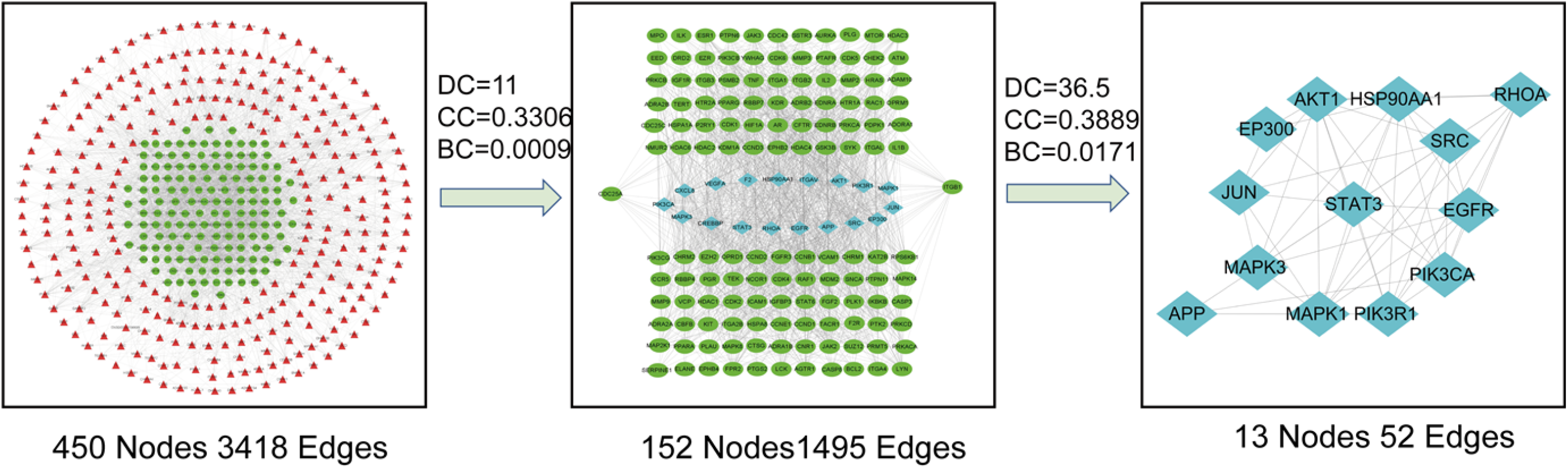
PPI network construction diagram of common targets.

#### 3.3.4. Enrichment analysis

The GO analysis of the corresponding targets of the active components of JFKG was carried out by using DAVID database. The screening condition was FDR < 0.001; and the biological process (BP); molecular function (MF) and cellular components (CC) were screened. The drawing was made by bioinformatics (Fig 8D1). Among them; the top one in BP analysis is signal transduction; protein phosphorylation; positive regulation of transcription from RNA polymerase II promoter; response to drug; negative regulation of apoptotic process; positive regulation of cell proliferation; negative regulation of transcription from RNA polymerase II promoter; proteolysis; protein autophosphorylation; cell proliferation; CC analysis; and the top one is plasma membrane; nucleus; cytoplasm; cytosol; nucleoplasm; extracellular exosome. The top MF is protein binding; binding; protein kinase activity; protein serine/threonine kinase activity; zinc ion binding; which speculates that JFKG in the treatment of CS may be the result of a complex multi-way synergistic effect.

**Fig 8.**
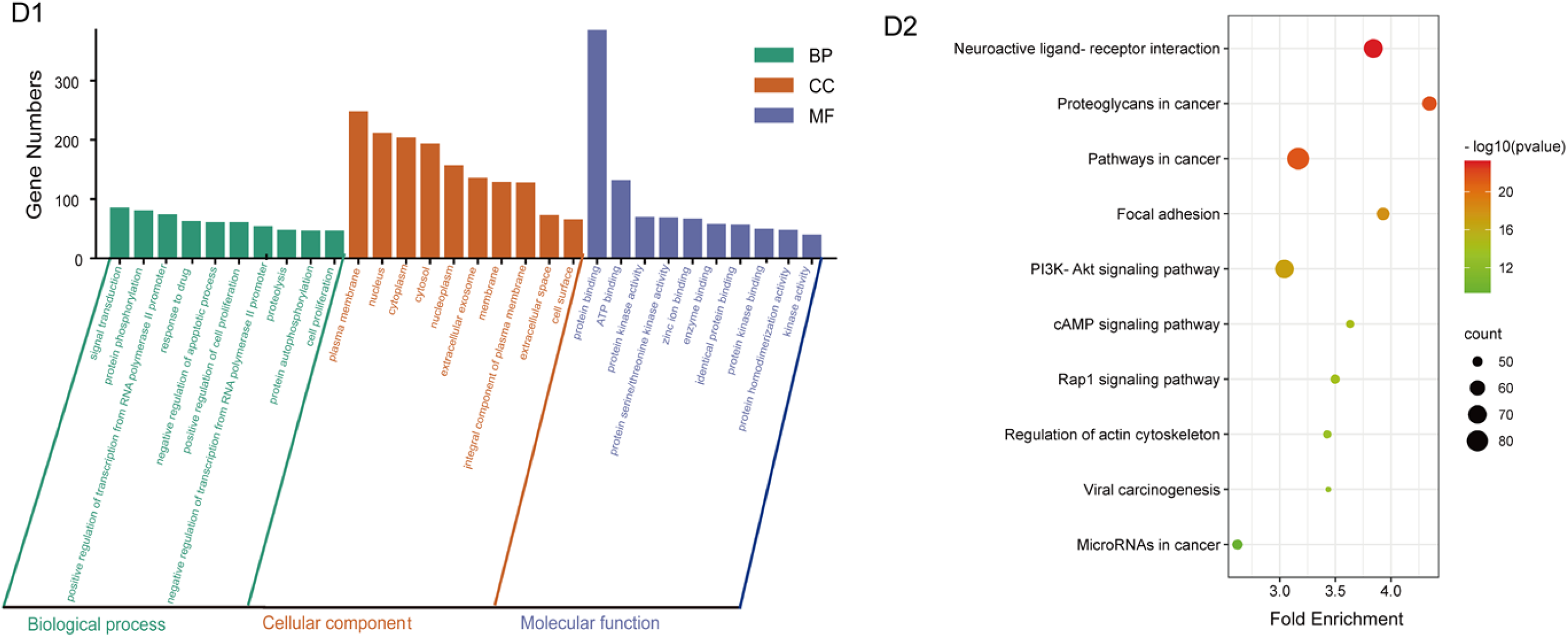
The GO enrichment analysis(D1) and the KEGG analysis(D2)

The results of KEGG signal pathway analysis: a total of 136 signal pathways (P<0.05) were identified and the first 10 pathways with many enriched genes were selected (Fig 8D2). The mechanism of JFKG in treating CS mainly involves Pathways in cancer; Neuroactive ligand-receptor interaction; PI3K-Akt signaling pathway; Proteoglycans in cancer; Focal adhesion; MicroRNAs in cancer; Rap1 signaling pathway; cAMP signaling pathway; Regulation of actin cytoskeleton; Viral carcinogenesis; and other pathways. Therefore; it is speculated that the treatment of CS with JFKG may be closely related to the above pathway.

#### 3.3.5. Construction of “TCM-component-target-pathway-disease” Network

To better clarify the relationship among components; targets; and pathways; we use Cytoscape3.8.2 to construct a “TCM-component-target-pathway-disease” network (Fig 9). Through this network; we can directly show the effective substances of JFKG in treating CS and its possible mechanism.

**Fig 9.**
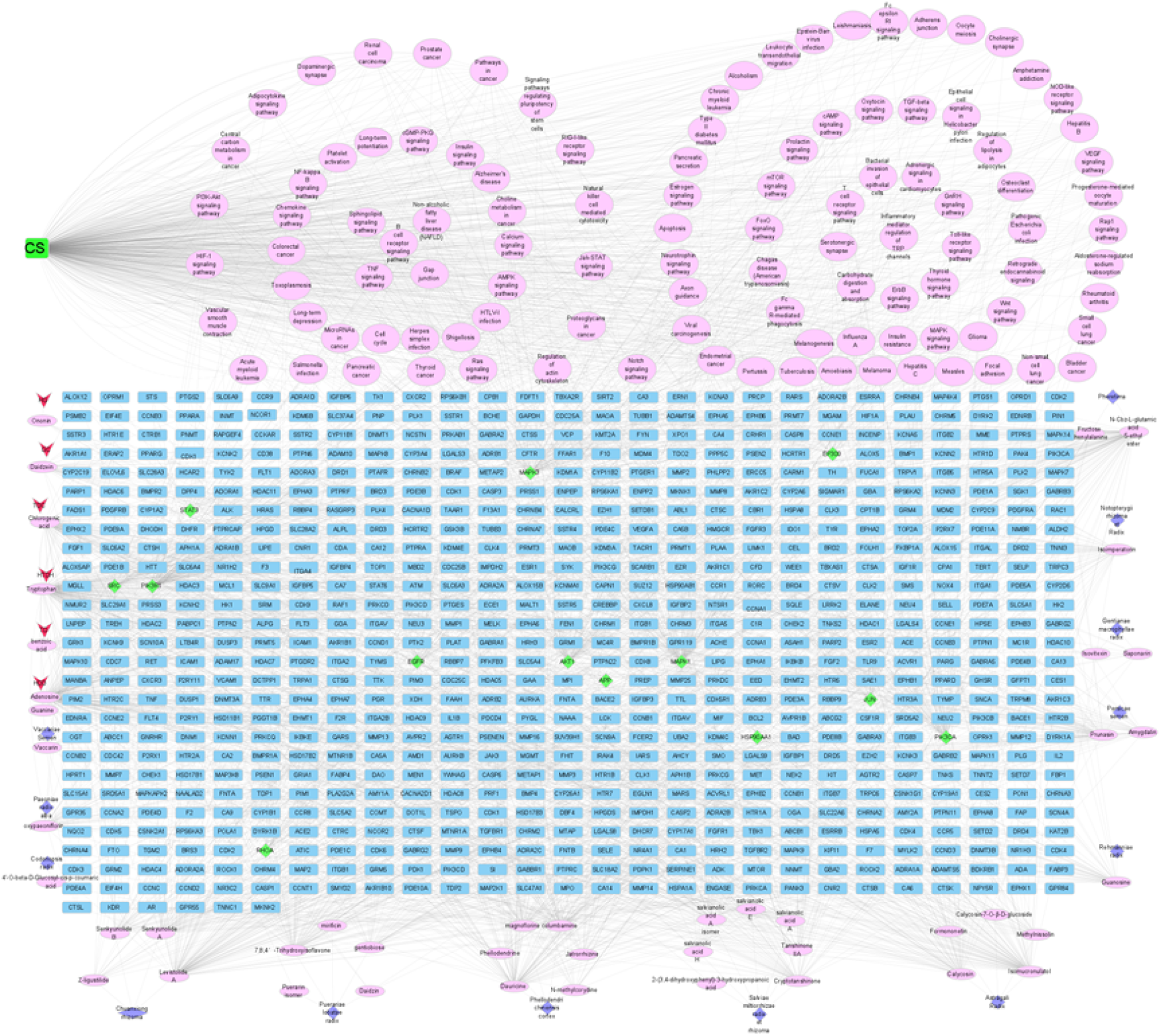
TCM-component-target-pathway-disease Network of JFKG.

## 4. Discussion

With the popularization of clinical traditional Chinese medicine; the phenomenon of too much flavor; large dose and the combination of similar drugs in the current prescription of traditional Chinese medicine is more prominent; which has aroused widespread concern [24]. The study of chemical composition and pharmacodynamics has become the main content of the basic research of traditional Chinese medicine formula granules; using the fingerprint of modern science and technology; LC-MS to detect specific substances and related techniques such as one test and multiple evaluation [25]. The establishment of a comprehensive and overall quality control index has become a key problem to be solved in front of the development of traditional Chinese medicine formula granules. What’s more; LC-MS has strong separation ability and structure identification characteristics. This experiment can preliminarily clarify the medicinal material using the high resolution and specificity of UPLC-QE-Orbitrap-MS. Thus; the evaluation and control of the quality of traditional Chinese medicine must truly reflect its “internal” and “comprehensive” quality [26].

This study; aims at exploring the pharmacodynamic material basis of JFKG and lay a foundation for the study of its mechanism of action. A feasible UPLC-QE-Orbitrap-MS method was established to identify the components of JFKG effective extraction in vivo and in vitro. 97 components were identified in vitro; including 21 flavonoids; 17 organic acids; 15 terpenoids; 8 alkaloids; and other compounds. After oral administrating JFKG effective extraction; in the plasma of SD rats; 16 prototype components were identified; including 4 flavonoids; 3 organic acids; 2 alkaloids; 2 terpenoids; 2 sugars; 1 coumarin and 2 other compounds; which mainly originate from Puerariae lobatae Radix; Salviae miltiorrhiza; Astragali Radix; Phellodendri chinensis Cortex and other traditional Chinese medicines. In fact; Salviae Miltiorrhizae Radix et Rhizoma has the effects of promoting blood circulation and removing meridian obstruction; eliminating blood stasis and relieving pain; studies showed that tanshinone IIA; salvianolic acid B and so on; theirs main active constituent; can lower blood lipids; reduce blood viscosity; dilate blood vessels; inhibit platelet aggregation and nuclear release; have an antithrombotic effect; improve microcirculation; and repair injured vascular endothelial cells [27–28]. Astragalosides in Astragali Radix have anti-inflammatory; analgesic and immunomodulatory effects; which may be related to reducing capillary permeability; inhibiting neutrophil migration; reducing PLA activity and reducing the production of proinflammatory factors [29].

Next; the identified JFKG chemical constituents in vitro were further researched by network pharmacology. It was found that there were 62 medicinal components and 2818 potential targets in JFKG and 6923 CS related disease targets; including 531 common targets between JFKG and CS. The enrichment analysis of GO biological process and KEGG signal pathway showed that JFKG could regulate 13 core targets (PIK3R1; PIK3CA; APP; SRC; MAPK1; MAPK3…); 96 related pathways (PI3K-Akt signaling pathway; cAMP signaling pathway; MAPK signaling pathway; TNF signaling pathway and other pathways…) and inhibit inflammatory reaction and regulate immune function to treat CS. According to the analysis of genetic interaction; PIK3R1 gene is considered to have a potential pain-related regulatory function [30]; and its expression level may be related to the pathogenesis of osteoarthritis [31]; which lays a foundation for the pharmacological effect of JFKG on promoting blood circulation and relieving pain. Pharmacological data in previous research showed that PI3K-Akt signal pathway can promote angiogenesis and promote blood flow by activating important vasoactive factors such as vascular endothelial growth factor and nitric oxide [32–33]. TNF signaling pathway can induce a variety of intracellular signal transduction; including apoptosis; cell survival; inflammation and immune Rothman [34]. It has been found that rat cervical dorsal root ganglion compression can cause pain; while TNF-α antibody can block the related signal transduction and increase the pain threshold. MAPK signaling pathway can regulate the survival and apoptosis of neurons and alleviate the symptoms of nerve injury caused by CS [35–36]. The most common second messenger is cyclic adenosine monophosphate; which is promoted by the activation of adenylate cyclase after the ligands of the G-protein-coupled receptors; such as hormones and neurotransmitters; are linked. Not only is the production and degradation rate of cyclic adenosine monophosphate sensitive to a variety of extracellular and intracellular signals; but changes in intracellular cyclic adenosine monophosphate level will also affect intracellular signal transduction pathways; thereby regulating protein activity; gene expression and cell function; and thereby regulating cell metabolism; growth; differentiation and apoptosis [37–38]. According to the report; the increase of cyclic adenosine monophosphate level in nerve tissue not only helping to reduce the secondary pathological reaction after spinal cord injury; but also promoting nerve regeneration [39]. Other signal pathways with high correlation have not been reported in the literature and need to be further explored to confirm their mechanism.

Traditional Chinese medicine is often treated by oral decoction; which determines that the therapeutic effect of traditional Chinese medicine is not produced until it is absorbed by the intestines and stomach or metabolized. Therefore; looking for the components that may have therapeutic effect after entering the blood is the key object to analyze the material basis of the efficacy of traditional Chinese medicine [40]. However; many ingredients in traditional Chinese medicine do not need to be absorbed into the blood to play their corresponding role; especially some preparations of traditional Chinese medicine; such as external application or liquid bath; can be effective in the treatment of skin diseases [41]. If we only look for the active ingredients of traditional Chinese medicine from the blood-entering ingredients; it is bound to ignore some non-blood-entering ingredients with strong pharmacological activity. Secondly; even at the maximum dose; the concentration of some components in serum is still too low; how to ensure the detection of these components is still a difficulty. In addition; in the current study; too much attention has been paid to the transitional components in the serum at the peak of blood concentration; ignoring the effective components before and after the peak [42]. So to clarify the effective components of traditional Chinese medicine more comprehensively and accurately; this study used the extracted components of JFKG to construct the “TCM-component-target-pathway-disease” network; but did not select the blood ingredient.

## 5. Conclusions

In this study; the effective components and blood components of JFKG were analyzed for the first time; and the effective components were used to construct a “TCM-component-target-pathway-disease” network. This study not only providing an efficient and rapid analysis method for the qualitative analysis and quality control of chemical components and further clarifying the medicinal material of JFKG but also promoting the development of traditional Chinese medicine and innovate national traditional Chinese medicine.

## Acknowledgements

The financial support was provided by the National Natural Science Foundation of China (NSFC) (grant number 81373942); the Key Science and Technology Research Projects of Tibet Autonomous Region of China (XZ201801-GA-16) and the Education Department Foundation of Liaoning Province (LZ2020074). We are very grateful for the support of Analysis and Testing Center of Beijing University of Chinese Medicine. We are also very grateful for the support of Analysis and Testing Center of Beijing University of Chinese Medicine.

